# Modeling and Inference Methods for Switching Regime-Dependent Dynamical Systems with Multiscale Neural Observations

**DOI:** 10.1101/2022.06.09.494416

**Authors:** Christian Y Song, Han-Lin Hsieh, Bijan Pesaran, Maryam M Shanechi

## Abstract

Realizing neurotechnologies that enable long-term neural recordings across multiple spatial-temporal scales during naturalistic behaviors requires new modeling and inference methods that can simultaneously address two challenges. First, the methods should aggregate information across all activity scales from multiple recording sources such as spiking and field potentials. Second, the methods should detect changes in the regimes of behavior and/or neural dynamics during naturalistic scenarios and long-term recordings. Prior regime detection methods are developed for a single scale of activity rather than multiscale activity, and prior multiscale methods have not considered regime switching and are for stationary cases. Here, we address both challenges by developing a Switching Multiscale Dynamical System model and the associated filtering and smoothing methods. This model describes the encoding of an unobserved brain state in multiscale spike-field activity. It also allows for regime-switching dynamics using an unobserved regime state that dictates the dynamical and encoding parameters at every time-step. We also design the associated switching multiscale inference methods that estimate both the unobserved regime and brain states from simultaneous spike-field activity. We validate the methods in both extensive numerical simulations and prefrontal spike-field data recorded in a monkey performing saccades for fluid rewards. We show that these methods can successfully combine the spiking and field potential observations to simultaneously track the regime and brain states accurately. Doing so, these methods lead to better state estimation compared with single-scale switching methods or stationary multiscale methods. These modeling and inference methods effectively incorporate both regime-detection and multiscale observations. As such, they could facilitate investigation of latent switching neural population dynamics and improve future brain-machine interfaces by enabling inference in naturalistic scenarios where regime-dependent multiscale activity and behavior arise.

## 1. Introduction

Linear dynamical systems (LDS) have served as the foundational models for brain-machine interfaces (BMIs) to control neural prosthetic devices and for the study of neural population dynamics underlying behavior [1–5]. An LDS assumes that an unobserved continuous-valued state related to behavior is linearly encoded in the neural observations and that this state evolves over time with time-invariant dynamics. We refer to this state as the brain state, which can either directly represent behavior parameters such as movement kinematics, or be a latent state that gives rise to neural activity and/or behavior [2,3,6]. Inference algorithms such as the Kalman filter then rely on these LDS model assumptions to efficiently estimate brain states from neural observations. However, these model assumptions become less appropriate as neurotechnologies measure multiple spatiotemporal scales of neural activity using simultaneous field potential and spiking modalities, and as they do so during more complex and naturalistic behaviors.

First, when both spiking and field potentials are being recorded, the linear observation model in LDS will need to be modified to accommodate for the statistical and time-scale differences of these modalities, with field potentials being continuous signals and spikes being binary action potential events with faster time-scales. We refer to the combination of both spiking and field potential signals with their different statistical and timescale profiles as multiscale observations. Second, for more naturalistic or complex behaviors, neural dynamics may change over time depending on the regime of behavior. We refer to periods of time with unique brain state dynamics and encoding as regimes, which we label at each time point by a discrete-valued variable termed the regime state. Thus, there is an unmet need to develop modeling and inference methods that not only use multiscale observations but also relax stationarity assumptions to include regime-dependent dynamics and encoding.

This need is motivated by the observation that behavior is encoded across multiple spatiotemporal scales of neural activity [5, 7–12]. It is also motivated by advances in neural devices that can simultaneously record spiking activity from individual neurons and field potentials – in the form of local field potentials (LFP) or electrocorticogram (ECoG) – that reflect the activity of larger neural populations [13]. Prior studies have developed modeling and filtering methods for multiscale spike-field observations called the multiscale dynamical system model and the multiscale filter, respectively [7,8, 14], which model the field features as a Gaussian process as in a traditional LDS model but also model the spiking activity as an additional point process that nonlinearly encodes the brain state. These studies found that spiking and LFP reflect a shared low-dimensional neural state that explains naturalistic behavior [8]. They also found that the multiscale filter improves movement decoding compared to both a Kalman filter and a point process filter [7], which both use only a single scale of activity in the form of field features or spikes respectively. Thus, explanatory power and decoding performance can improve by developing a model that can effectively combine information from multiple neural activity scales. However, these studies also assumed stationarity over time. Thus, a major question is how to extend these multiscale methods to allow for non-stationarity, which is especially critical in more naturalistic setups.

One way to relax the stationarity assumption is to develop adaptive methods, which has been done for single-scale state-space models in BMIs (e.g., [15–22]). Adaptive methods are suitable for cases where the model might slowly change over time due to for example neural plasticity or motor learning but not for all cases where the stationarity assumption must be relaxed. Specifically, sometimes the model could abruptly switch parameters due to a regime change, where a regime can represent, for example, the stage of a task, or an internal state such as stress, engagement, or attention [23–26]. For the case of single-scale LDS, prior works in neural engineering and computer vision [27, 28] have handled cases with a switching regime-dependent nature using the switching linear dynamical system (switching LDS) model. A switching LDS augments an LDS by assuming that there exists an additional governing regime state dictating which parameters to use based on which regime is in power. The regime state can then switch between a discrete finite set of regimes with Markovian dynamics. However, this extension has not been done for cases with multiscale observations that do not fit within the LDS framework and remains a challenge.

Figure 1 depicts the appropriate use case for a unified model where the behavior-related brain state is encoded in multiscale observations with an overarching switching regime state that governs the dynamical and encoding parameters. Current inference methods for switching LDS can only handle a single scale of observations [29–31], while current methods for multiscale dynamical systems assume the model is stationary [8, 14]. Thus, novel inference algorithms must be developed to estimate both the behavior-related brain state and the regime state given simultaneous continuous-valued Gaussian observations and binary-valued point process observations.

**Figure 1.**
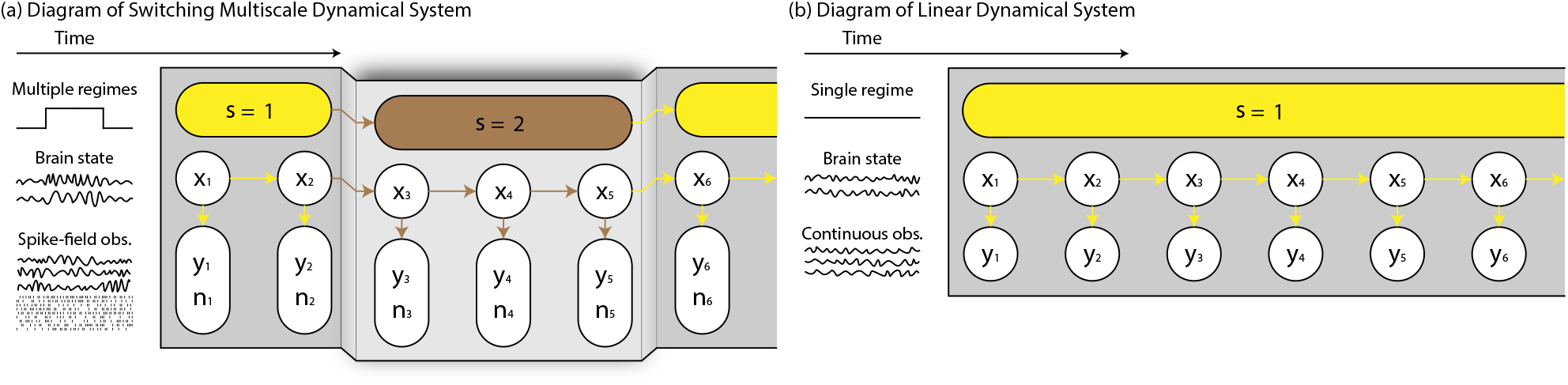
Visualization of models of dynamical systems. (a) Model of a Switching Multiscale Dynamical System (SMDS) in which the regime in power s dictates both the dynamics of the brain state **x** and the encoding in multiscale spike-field observations **y** and **n**. (b) Model of a Linear Dynamical System (LDS) with stationary brain state dynamics and encoding in only a single continuous observation scale.

Here, we develop the modeling and inference methods for the switching multiscale dynamical system (SMDS) and demonstrate them in both extensive numerical simulations and monkey spike-LFP activity during a saccadic eye movement task with various regimes. The SMDS model extends the LDS model by appending an additional binary-valued point process observation modality and an additional discrete-valued regime state, which dictates the dynamical and encoding parameters. The brain state is encoded nonlinearly in the instantaneous firing rate of the point process and linearly in the Gaussian process while the regime state evolves as a first order Markov chain. We also derive both a filter and a smoother for the SMDS model. The filter termed the Switching Multiscale Filter (SMF) is causal and computationally tractable and thus practical to use in real-time decoding and BMI applications. The smoother termed the Switching Multiscale Smoother (SMS) can leverage the entire length of data to reliably perform non-causal and more accurate estimation when real-time processing is not needed.

We first validate the developed methods with extensive Monte Carlo spike-field simulations and show their success and robustness in estimating both brain and regime states across diverse settings and parameter ranges. Specifically, SMF and SMS more accurately estimate the brain and regime states compared to multiscale inference methods that use the same spike-field activity but assume stationarity. Also, the new methods successfully combine information from multiple neural observation scales to better estimate the brain and regime states compared to when using a single scale of observation. Finally, the SMS smoother reliably yields more accurate estimation of both brain and regime states compared to the SMF filter across every single parameter setting tested. Indeed, we show that even in the single-scale case, prior smoothing methods do not generalize as well as SMS across system settings. We then show the success of the SMF method in multiscale spike-LFP data recorded from a monkey performing saccades for fluid rewards. Specifically, SMF can detect different regimes in multiscale spike-LFP dynamics that are associated with different task stages and successfully combines information from spiking activity and LFP features to improve estimation of regime and brain states. Finally, the decoded brain states can accurately classify the direction of the executed saccade.

## 2. Methods

In this section, we present the modeling and inference methods for the SMDS model with an illustration of the developed inference methods in Figure 2. We then set up the validation methods consisting of extensive Monte Carlo simulations of SMDS models and applications of our methods in data collected from a non-human primate (NHP). We denote column vector values with lowercase bold letters, matrices with uppercase bold letters, continuous-valued densities with *f* (·), and discrete-valued distributions with *P*(·).

**Figure 2.**
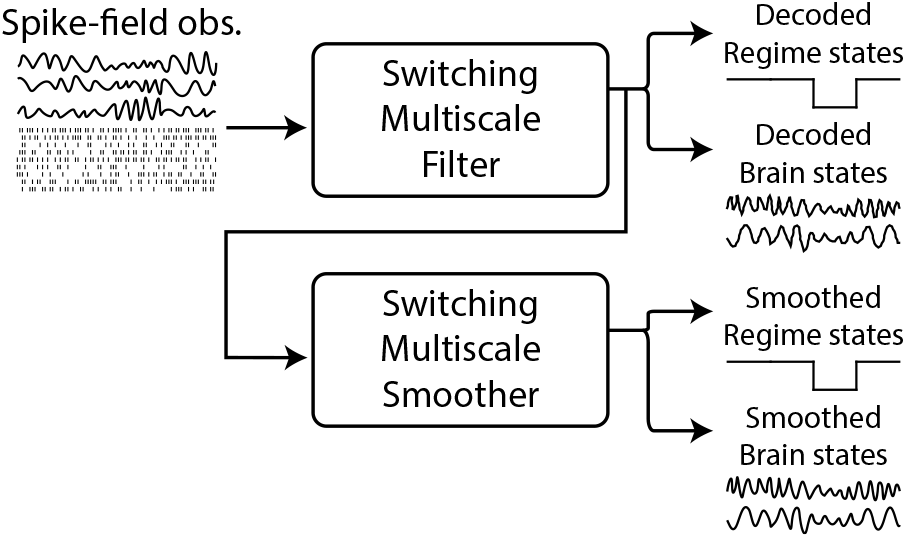
Illustration of inference methods for SMDS models. The switching multiscale filter (SMF) takes spike-field activity to causally decode unobserved brain and regime states. The values calculated in SMF can then be used in the switching multiscale smoother (SMS) to further smooth estimates of brain and regime states non-causally using future spike-field activity.

### 2.1. Switching multiscale dynamical system (SMDS)

We first detail the switching multiscale dynamical system (SMDS) model. The regime state *s_t_* is a discrete random variable that can take one of *M* possible values from 1 to *M* and describes the current regime of the system. For example, this regime state can represent the current stage within a multistage task. This regime state evolves with first order Markovian dynamics as:

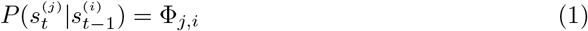

where **Φ** is the transition matrix, and 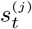 and 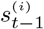 respectively denote *s_t_* = *j* and *s*_*t*–1_ = *i*. The diagonal elements Φ_*j,j*_ describe the probability of staying in the same regime *j* while the off-diagonal elements Φ_*j,i*_ describe the probability of transitioning from regime *i* to *j* at every time evolution. We denote the initial distribution of the regime state s_1_ as ***π***. The regime state then dictates the parameters of the dynamical model at every point in time.

The brain state represents the state of neural activity that relates to the behavior being performed, e.g. eye movement direction or arm movement kinematics. We model the brain state **x**_*t*_ as a random walk driven by zero mean Gaussian noise **w**_*t*_ with covariance **Q**(*s_t_*):

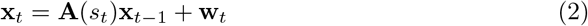

The dynamics **A**(*s_t_*) can take on one of *M* possibilities {**A**^(1)^,…**A**^(*M*)^} where 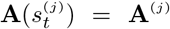. The same notation holds for the noise covariance **Q**(*s_t_*). The initial brain state **x**_0_ is assumed to be Gaussian with mean ***μ***_0_ and covariance **Λ**_0_. The above constitutes the brain state transition model. We refer to the parameters associated with the brain state dynamics in (2) as the dynamics equation parameters.

We model the encoding of **x**_*t*_ in simultaneously recorded field potential features and spiking activity. Prior work has shown that behavior can be linearly mapped to the log-power of various frequency bands of field potential activity [6,8–10,32–38]. Thus, we model the field features **y**_*t*_ as:

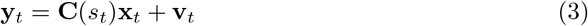

where **v**_*t*_ is Gaussian noise independent of brain state noise **w**_*t*_ with zero mean and covariance **R**(*s_t_*). The parameters **C**(*s_t_*) and **R**(*s_t_*) are chosen from *M* possibilities by the regime state.

The spiking activity for neuron *c* is then modeled as a point process with **x**_*t*_ encoded in the instantaneous firing rate λ_*c*_ [7,8,14,17,18,39–42]. We denote the time-step by Δ and choose it small enough to contain at most one spike, thus generating a binary-valued signal indicating the presence or absence of a spike. The point process likelihood distribution of the spiking activity from *C* neurons 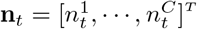 is:

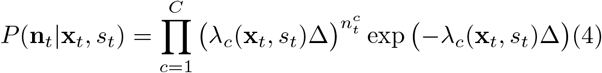

with the firing rate in the form of an exponential function:

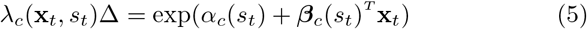

We also assume that the activity of the neurons are conditionally independent of each other conditioned on the brain state **x**_*t*_ [17,39,40,42]. The parameters *α_c_*(*s_t_*) and *β_c_*(*s_t_*) are each chosen from *M* possibilities by the regime state. We refer to the parameters related to observation encoding in (3) and (5) as the observation equation parameters.

We can now combine the two observation modalities into a multiscale observation modality with a joint likelihood density, which is a key feature of the SMDS model. Prior studies have shown that a reasonable assumption is that the field features and spiking activity are conditionally independent given **x**_*t*_ [7,8]. So, we can write the joint likelihood density as

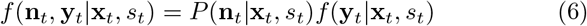

The SMDS model also allows for the possibility that the time-scale of field features may be longer than that of the recorded spiking activity, meaning that the former may have a longer time-period between their consecutive observation samples. In this case, since we have to set the time-step to be the shorter time-period for consecutive spike events, there exist time-steps with no field observation. We thus model unobserved field features by setting **C** = 0 or **R** = ∞. Finally, we denote the multiscale spike-field observation at time *t* as **h**_*t*_ = [**y**_*t*_, **n**_*t*_]^T^. Together, (1) – (6) constitute the SMDS model. We highlight that this model extends both the switching LDS and the multiscale dynamical system by accommodating not only multiscale observations but also regime-dependent non-stationarity.

### 2.2. Inference algorithms

We develop inference algorithms for the SMDS model that aim to obtain the minimum mean squared error (MMSE) estimate of the brain and regime states given available multiscale observations. The MMSE state estimator based on observations up to time τ is known to be the mean of the posterior density [30], and thus we aim to find the following terms for the brain and regime state estimation, respectively:

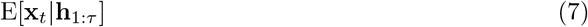

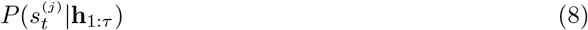

where *τ* is equal to the present time *t* for filters and equal to some future time *T* for smoothers.

Computing the above terms exactly, however, requires an exponentially increasing computational cost as time grows. To see this, focusing on the decoding problem for (7), we can expand it to consider each potential regime state sequence:

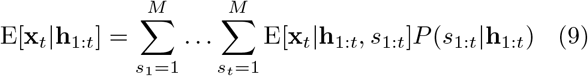

In the inner expectation term E[**x**_*t*_|**h**_1:*t*_, *s*_1:*t*_], given each potential sι_:t_ sequence, the regime is known for each time t. So finding the inner expected brain state term is a simple filtering problem. However, the number of potential sequences and thus the number of computations grows exponentially with time, where at time *t* with *M* possible regimes, *M^t^* sequences must be considered. Thus, the inference algorithms that we develop must approximate the above terms to reconcile this challenge of intractability and ensure practicality.

#### 2.2.1. Switching multiscale filter

We first formulate the overall recursive approach that the switching multiscale filter takes and introduce the following notation:

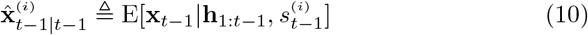

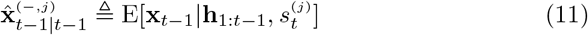

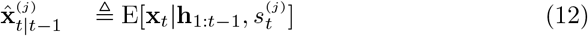

Through the rest of this work, *i* will be used to index the regime state at time *t* – 1 and *j* for time *t*.

We start by expanding the posterior density of the brain state over the potential regimes at time *t* – 1:

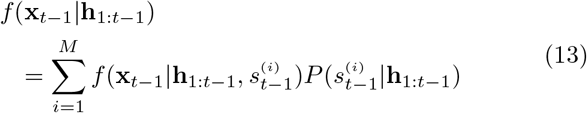

where the mean of (13) would be the MMSE estimate of the brain state at time *t* – 1. We assume that each 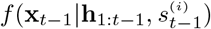 a Gaussian with mean and covariance 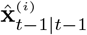 and 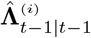, respectively. Thus, (13) can be viewed as a mixture of Gaussians whose density is completely described by the below sufficient statistics comprised of the mean and covariance of each Gaussian and its relative weight 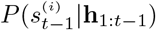):

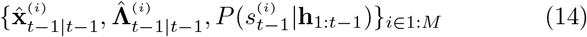

This weight is also critical in finding the estimate of the regime state. We then perform the same expansion of the brain state density at time *t*:

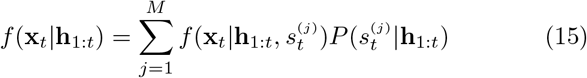

We also assume that each 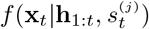 is also a Gaussian and that (15) is a mixture of Gaussians with sufficient statistics:

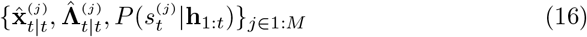

The inference goal is to compute (16) from (14) recursively. The approach of SMF then starts each recursion with the terms (14) being known and incorporates the new multiscale observations at time *t*, i.e. **h**_*t*_, to calculate the next set of terms (16) to complete the recursion. A summary of SMF can be seen in Algorithm 1, which we now derive.

#### Algorithm 1 Switching Multiscale Filter

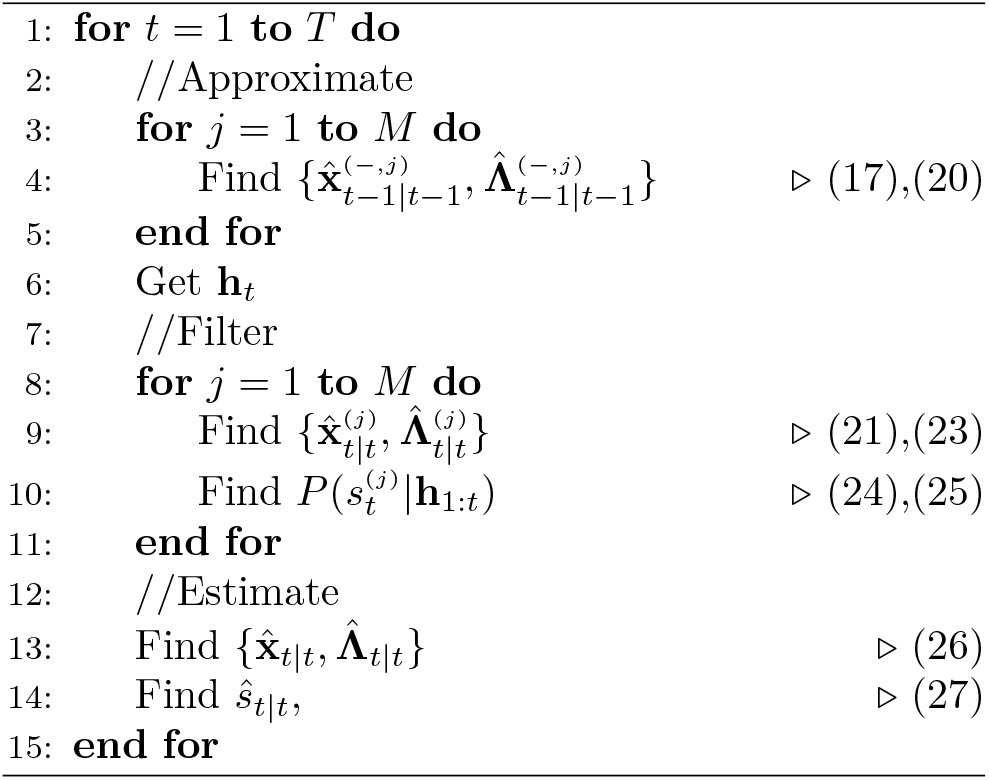

To begin the derivation of SMF, we first increment the regime state for each 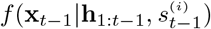 such that it is conditioned on 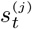 instead of the current 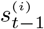 to yield the density 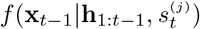. This is done through marginalization and Markov conditional independence properties:

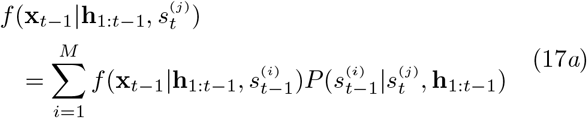

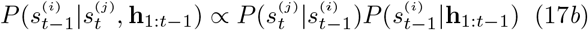

We can see that (17*a*) is a mixture of Gaussians because we assumed 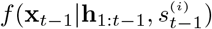 is a Gaussian in (13) and that the weights of the mixture (17*b*) can be calculated from (1) and (14). To ensure tractability, we approximate this mixture of Gaussians as a single Gaussian whose mean and covariance equal those of the mixture.

The overall mean and covariance, *μ* and **A**, of a general mixture of *M* Gaussians with individual means, covariances, and weights denoted by {**μ**^(*i*)^, **A**^(*i*)^, *p_i_*}_*i*∈[1,*m*]_ can be found through [43]:

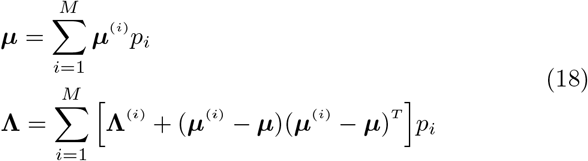

We will denote this process of finding the first and second moments of a mixture of Gaussians as:

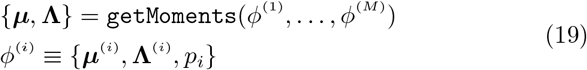

Thus, we approximate (17*a*) with a single Gaussian with mean and covariance:

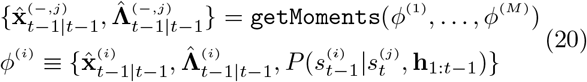

This is repeated for each regime *j* ∈ [1, *M*].

We now incorporate the next observation **h**_*t*_ and update the brain state density from 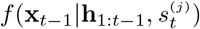) to 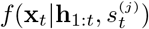 for each regime *j*. Because the regime is fixed at *j* for the above two densities, we do this by adapting a recently developed filter for stationary multiscale observations [7] due to the multiscale nature of **h**_*t*_ and using model parameters associated with regime *j*. The first step involves a prediction of the brain state to get the prediction density 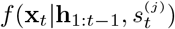 with mean and covariance:

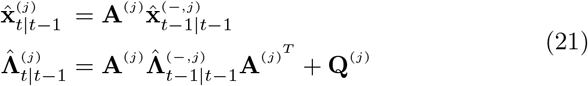

We then update the brain state to 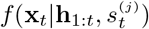 by expanding the density with Bayes rule:

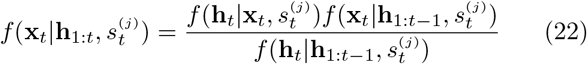

and approximating the density as a Gaussian with Laplace’s approximation with mean and covariance:

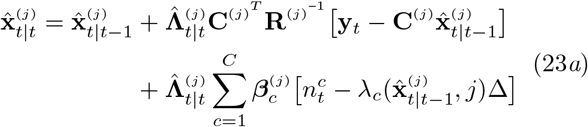

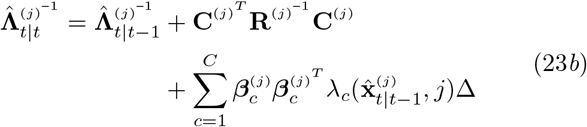

We note that without first introducing 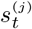 dependence while removing 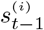 dependence from 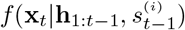 in (17) and (20), this filtering step would result in a brain state density dependent on both 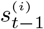 and 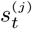 and thus would require additional steps to later remove the 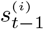 dependence. However, here, the above steps result in the desired mean and covariance terms in (16) without such a dependence.

We now focus on calculating the regime state distribution 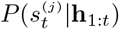. We first apply Bayes rule to the distribution:

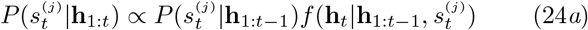

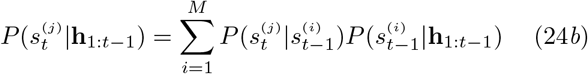

where (24b) (the first term in (24a)) is found from (1) and (14). However, the second term in (24*a*) is difficult to find due to the multiscale nature of the observations in the SMDS model. To overcome this challenge, we recognize that this second term is also the partition function (i.e. denominator) of the update density (22). Thus, we solve for it using (22) in terms of the prediction and update densities, which were both approximated as Gaussians in the Laplace approximation with means and covariances given by (21) and (23). We then evaluate this second term at 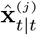 with some additional basic algebraic operations to derive the following solution:

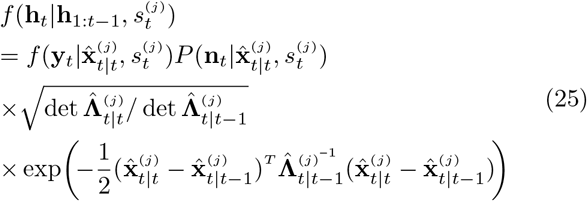

We use (25) with (24*b*) to calculate (24*a*) for each regime *j* to obtain the remaining terms in (16) and complete the recursion.

Finally, we calculate the MMSE estimate of the brain state *f*(**x**_*t*_|**h**_1:*t*_) as in (7) by obtaining the mean of the mixture (15):

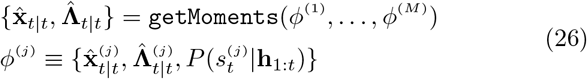

where *ϕ*^(*j*)^ is already available from above. The regime state estimate is then set to the regime with the largest posterior:

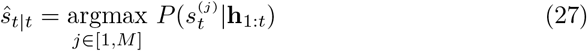

Together, (17) – (27) detail the SMF algorithm. The architecture for this method was inspired by a decoding algorithm for switching linear dynamical systems (LDS) called the interacting multiple model (IMM) [29, 30] algorithm. However, this IMM method only accepts Gaussian observations and thus is not suited for the SMDS model with multiscale observations.

#### 2.2.2. Switching multiscale smoother

We now present a fixed-interval smoother for the SMDS model termed the switching multiscale smoother (SMS). We run this method after SMF to obtain estimates of the brain and regime states at time *t* as in (7) and (8) given observations up to a fixed future time *T* ≥ *t*.

Compared to derivations of prior smoothing methods for switching LDS systems for single-scale observations, SMS introduces a key additional adjustment as described below. These prior smoothing methods for switching LDS in [31], which we call Kim’s method, and in [44], which is called Expectation Correction (EC), do not directly require any of the observations – whether multiscale or not. Instead, they rely only on the sufficient statistics computed by a filter like SMF and the assumption that (15) is a mixture of Gaussians. However, we find that directly applying these methods to the SMDS models or even to single-scale models (whether point process or Gaussian) leads to inconsistent/unreliable results across different system settings, with the prior smoothers doing worse than even filtering methods in some settings (see Results Section 3.3). Thus, we introduce a key additional adjustment in SMS which empirically yields a more reliable method that generalizes well across diverse settings (Results Section 3.3).

We introduce the overall approach of SMS first with the following notation:

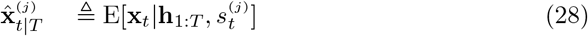

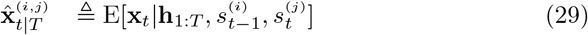

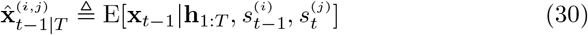

We begin by expanding the density of the brain state at time *t*:

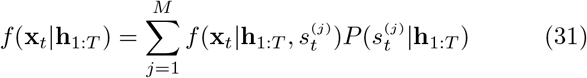

where the mean of (31) would be the MMSE estimate of the brain state given observations up to a fixed future time *T*. Similar to (13), we assume each 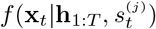 is a Gaussian with mean and covariance 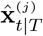 and 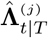 respectively, and thus we can view (31) as a mixture of Gaussians with sufficient statistics:

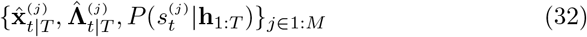

We then repeat the expansion on the brain state density at time *t*–1 with similar Gaussian assumptions, whose density and sufficient statistics are:

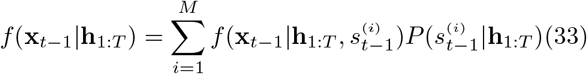

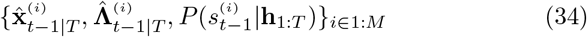

The inference goal is then to take the set of sufficient statistic terms in (32) and calculate the next set in (34) to complete the recursion. We note that the set (32) is calculated in the final recursion of SMF when *t* reaches *T*. Thus, after SMF is run in the forward time direction, SMS can then be run noncausally in the backwards time direction to yield more accurate estimates of the brain and regime states.

We start the derivation of SMS by focusing on the brain state. With the ultimate goal of finding the mean and covariance of 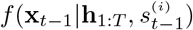 as in (34), we begin by finding the mean and covariance of an intermediate density 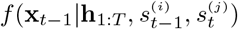 such that we can later remove the 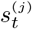 dependence. We first follow the derivation to the Rauch-Tung-Striebel (RTS) smoother and marginalize the intermediate density over **x**_*t*_:

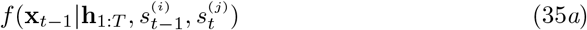

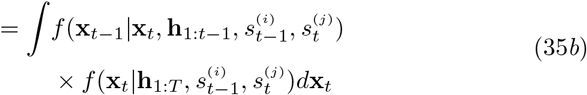

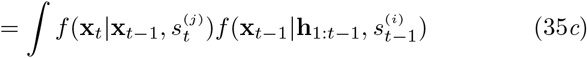

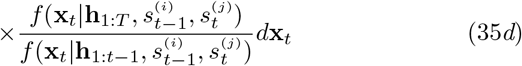

where (35c) and (35d) come from applying Bayes rule to the first density in (35b). As in the RTS smoother, the density’s mean and covariance can then be found with multivariate Gaussian properties [45]:

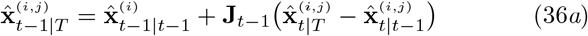

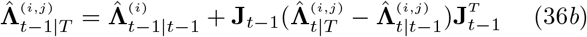

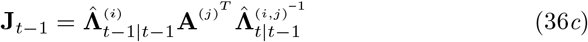

where, in (35c), **A**^(*j*)^ is the dynamics matrix associated with 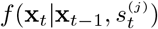, and 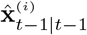 and 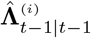 are the moments of 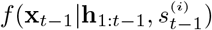. For (35d), 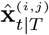 and 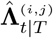 would be the mean and covariance of the smoothed density 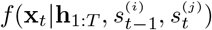, in the numerator while 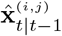 and 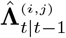 would be the moments of the prediction density 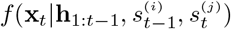 in the denominator. However, the challenge associated with these equations is that the moments for the smoothed density 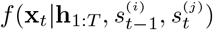 in (35d) are not actually known nor trivially computable, so we make the following assumption:

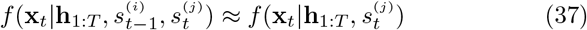

where the right density is known from (31) with mean and covariance 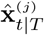 and 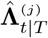 as in (32), and use these moments to replace 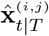 and 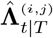. This assumption is also used in Kim’s method and EC and allows the mean and covariance of the intermediate density to now be computable.

However, we find that with certain model parameters, using these calculations as is can lead to poor estimation performance regardless of observation modality – whether single-scale or multiscale – as detailed in Section 3.3, and this motivated our key new additional adjustment. We hypothesized the problem lies in a discrepancy in (35*d*) between the prediction density 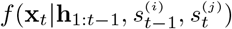 having an 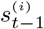 dependence while the smoothed density 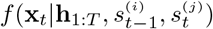 having its 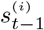 dependence assumed away in (37). Thus, to remedy this discrepancy, we make the prediction and smoothed densities consistent by making an additional equivalent assumption for the prediction density:

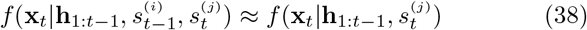

and replace the moment terms 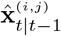 and 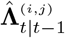 with those of 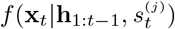, i.e. 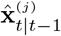 and 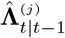, which we find in (21). The final equations for finding the mean and covariance of the intermediate density 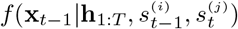 for SMS is then:

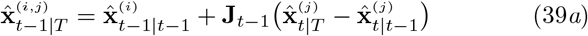

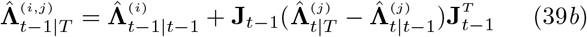

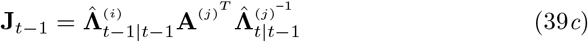

Thus, all ^(*i,j*)^ terms on the right hand side of (36) with 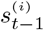 and 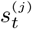 dependence become ^(*j*)^ terms with only 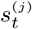 dependence. We show in Section 3.3 that this adjustment allows SMS to reliably smooth decoded estimates in cases where prior methods (Kim’s method and EC) could perform worse than even a causal filter.

We then continue the derivation by changing focus to the regime state. With the goal of finding 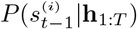 as in (34), we first find the joint distribution 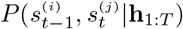:

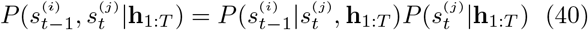

and make the following assumption to simplify computation:

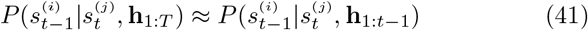

with the term on the right found in (17*b*). We repeat this for each (*i, j*) pair.

Finally, with 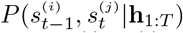 in (41) and the moments of 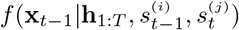 in (39) known, we remove their 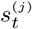 dependence to find the terms in (33) and ultimately the sufficient statistics (34) to complete the recursion. For the regime state, we marginalize the distribution over 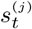:

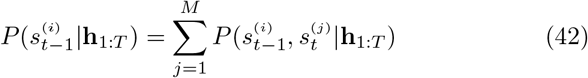

We similarly marginalize the brain state over 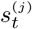:

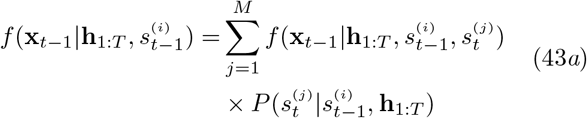

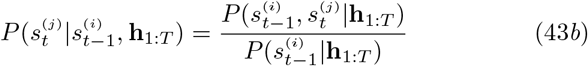

We assume 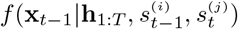 is Gaussian with moments 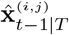 and 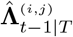 from (39). Thus, (43a) is a mixture of Gaussians which we approximate with a single Gaussian with the same mean and covariance 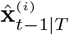 and 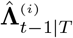:

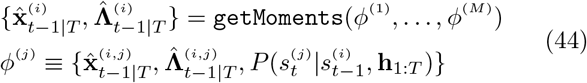

where the terms in *ϕ*^(*j*)^ are found from (39a), (39b), and (43b). We repeat this for each regime *i* ∈ [1, *M*] to get all terms in (34) from (42) and (44) and thus complete the recursion.

Finally, we calculate the MMSE estimate of the brain state density *f*(**x**_*t*–1_|**h**_1:*T*_) in (33) as:

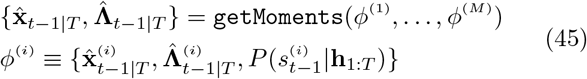

where *ϕ*^(*i*)^ are found from (44) and (42), and estimate the regime state as that with the largest posterior:

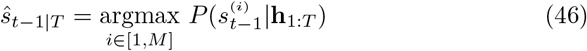

A summary of the algorithm can be seen in Algorithm 2. The main difference between SMS and prior smoothing methods is in the new additional assumption made in (38), which as we show in Section 3.3 is critical to enable reliable performance and generalizability under diverse system settings. We note that EC makes a different assumption in the joint regime state calculation in (41) with details in [44].

#### Algorithm 2 Switching Multiscale Smoother

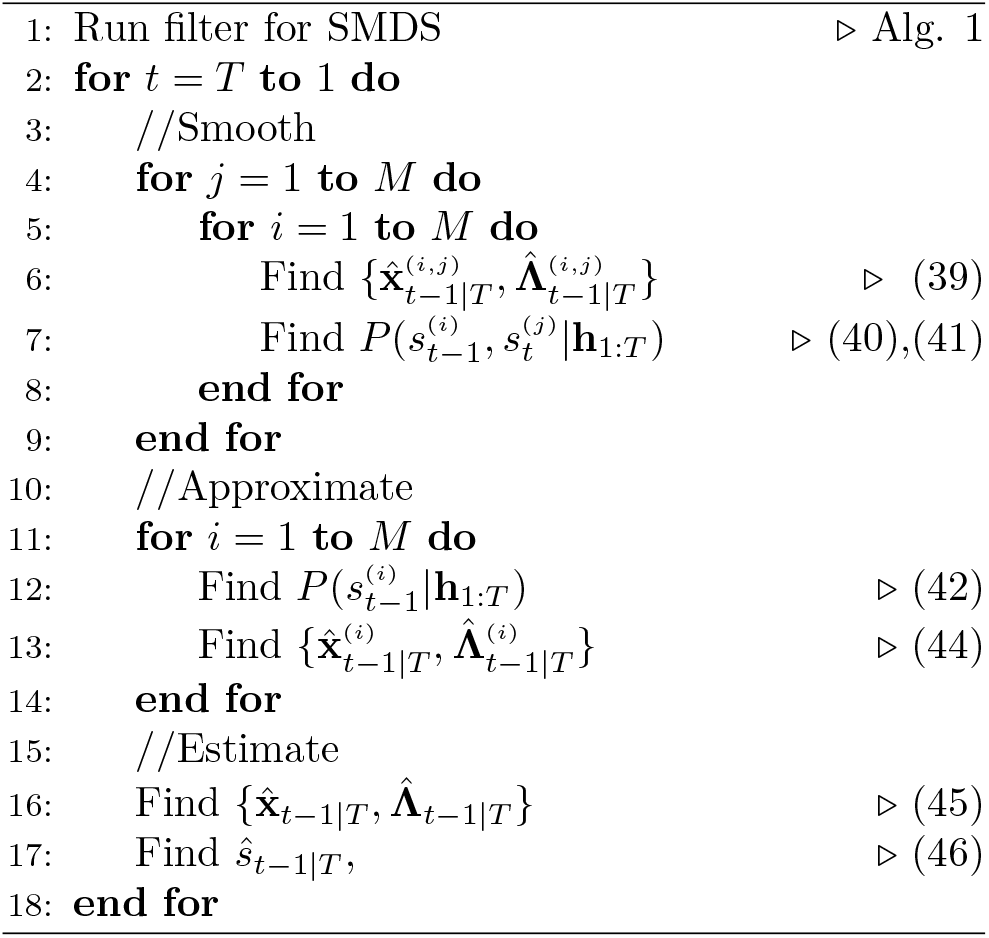

### 2.3. Performance Metrics

In simulations where the ground truth brain state is known, we compare the estimated brain states to the true brain states using the normalized root meansquare error (NRMSE) metric [14]:

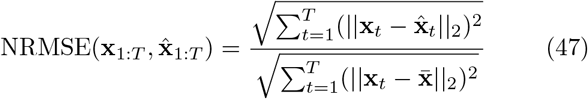

where **x**_1:*T*_ and 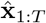 are the true brain states and estimated brain states respectively, || · ||_2_ is the Euclidean norm, and 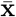 is the mean of the true brain states. This metric reflects more accurate estimation the lower it is, with perfect estimation at 0 and a chance level of 1, and with values above 1 representing worse than chance performance. We evaluate the estimated regime states based on accuracy, which is found from the proportion of estimated regime states that match the true regime states:

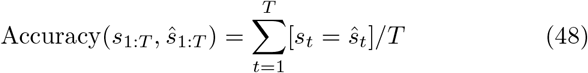

where 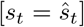 is 1 if the values match and 0 otherwise. Accuracy ranges from 0 to 1 and has a chance level of 1/*M* where *M* is the number of regimes.

### 2.4. Numerical simulation framework

We validate the developed inference algorithms using extensive Monte-Carlo numerical simulations. We simulate spike-field activity from randomly generated SMDS models under various simulation settings. We then measure the performance of the inference algorithms on the simulated data with the metrics in Section 2.3 and perform comparisons to prior methods, which either assume stationarity or allow switching but only use a single scale of observations. We ensure fair comparison of stationary and switching methods by estimating the parameters of both the stationary models and the switching models using maximum likelihood techniques [46]. For stationary methods, we fit a single set of parameters based on the entire training data, and for switching methods we fit the model parameters for each regime separately.

#### 2.4.1. Numerical simulation settings

For numerical simulations, we randomly form SMDS models under diverse simulation settings that we sweep. Details on specific values can be found in the Appendix and important parameters we sweep are summarized below. Across simulations, we assume the time bin Δ is 2 ms, and for each simulation setting, we randomly form 40 separate systems and simulate 40 × *M* seconds of data where *M* is the true number of regimes.

For the regime state settings, we sweep the number of regimes *M* and the dwell time *t_dwell_*, which is the average time spent within the same regime before a switch occurs. The dwell time can be calculated for regime *j* by *t_dwell_* = Δ/(1 – Φ_*j,j*_) where *Φ_j,j_* is from (1). We then assume that the probability of switching out of regime *j* to *i* is equal for all *i* ≠ *j* and simulate regime state realizations using (1).

For the dynamics equation parameters, we sweep the dimension of **x**_*t*_: dim(**x**_*t*_) = *d* and randomly form the dynamics matrix **A**^(*j*)^ and the noise covariance matrix **Q**^(*j*)^ per regime *j*. We repeat the below operations for each regime *j*, so the notation ^(*j*)^ will be omitted for clarity. Motivated by [8,14], we assume the eigenvalues of **A** are stable and comprised of *d*/2 complex-conjugate pairs {*r_i_e*^±*j*θ_*i*_^}_*i*∈[1,*d*/2_] as *r_i_* and *θ_i_* characterize the steady-state time dynamics of **x**_*t*_. The term *r_i_* dictates how fast the signal decays; for a purely real eigenvalue, a signal decays exponentially with a half-life of 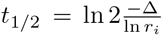. The term *θ_i_* dictates the frequency with which the signal oscillates, given by 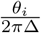 time units. In our simulations, we randomly select eigenvalues with decay half-lives between [27, 277] ms (i.e. a range of [0.95,0.995] for *r_i_*) and frequencies between [0.8,5] Hz (i.e. a range of [0.010, 0.063] for *θ_i_*) based on values seen in prior literature [4,8].

We then randomly select eigenvectors for each eigenvalue such that the dynamics matrix **A** can be decomposed into **A** = **UDU**^−1^ where **D** is a diagonal matrix of our selected eigenvalues and the columns of **U** are their corresponding eigenvectors. This is also known as the eigendecomposition of **A**. We call the collection of eigenvectors **U** for a given regime the eigenbasis and refer to the collection of **U** matrices across regimes as the eigenbases of the system. Unless otherwise stated, we assume the eigenbases are the same across regimes (i.e. stationary) to minimize transient signals that arise when regimes switch as the transients can interact with other settings and decrease interpretability of results. In some cases though, we also let the eigenbases switch to explore this effect; if we specify eigenbases can switch, we form **A** matrices such that the below holds to keep transient effects bounded:

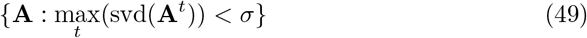

where svd gives the singular values of the input matrix and *σ* is what we call a transient control value that we choose ad hoc. We also randomly form the noise covariance **Q** with eigenvalues randomly chosen between 0.01 and 0.04. Finally, we simulate the brain states using (2) and the previously generated regime states.

For the observation equation parameters, we sweep various settings related to the simulated spike-field observations including the field feature sampling frequency and signal-to-noise ratio (SNR), and the maximum firing rate of the spiking activity with ranges that are common in neural data [7, 8]. For the field features, we assume the number of features to be 60 and randomly select parameters per regime based on the swept SNR to generate the observations using (3). For the spiking activity, we assume the number of neurons to also be 60 and randomly choose a base firing rate between 3-5 Hz. We then randomly select the parameters based on the base and maximum firing rates and generate observations using (5).

### 2.5. Experimental data

To demonstrate how our method performs on regime classification by fusing information from multiscale observations in neurophysiological data, we apply it to spiking and local field potential (LFP) activity in the lateral prefrontal cortex (lPFC) of one male rhesus macaque monkey (*Macaca mulatta*) performing saccadic eye movements for fluid rewards. All surgical and experimental procedures were performed in compliance with the National Institute of Health Guide for Care and Use of Laboratory Animals and were approved by the New York University Institutional Animal Care and Use Committee.

Each trial started with the monkey fixating on a central fixation point for a Baseline interval (400-800 ms). Following the Baseline interval, the central fixation point was extinguished and two response targets were illuminated at random locations on a circle separated by at least 60 degrees. Response targets were centered on fixation with eccentricity 10 degrees of visual angle. Following response target onset, which served as a Go cue, the monkey executed a saccade to one of the two targets for a fluid reward. In the following, we discretize the direction of the saccadic response by sub-dividing the response target circle into eight equal-sized bins as seen in Figure 3. The Start, End, and Target Onset/Go task events are defined as the trial start time, trial end time, and target onset time, respectively (Figure 3). Thus, each trial has two regimes, corresponding to the time periods before and after response target onset and referred to as Start-Go and Go-End regimes, respectively. The average trial length was 1.36 seconds, with the average time from the start of a trial to the Go cue being 0.697 seconds, the average time from the Go cue to the saccadic response being 0.176 seconds, and the average time from the Go cue to the end of a trial being 0.668 seconds.

**Figure 3.**
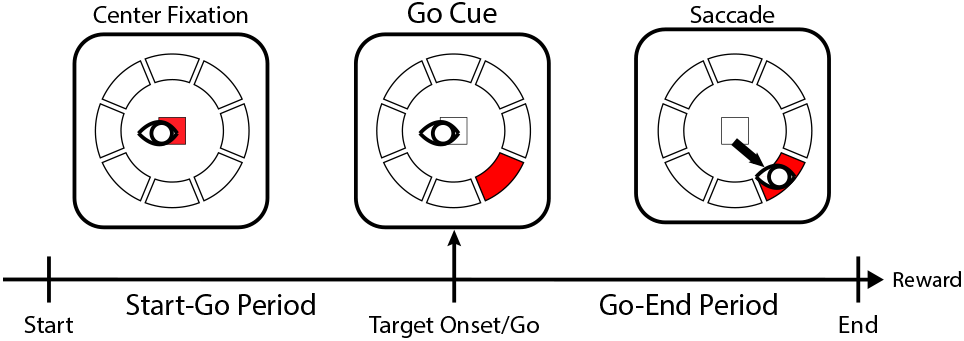
Behavioral task overview. A monkey performed a behavioral task to earn fluid rewards by executing a saccadic eye movement response. The direction of the saccadic response was discretized by sub-dividing the response target circle into eight equal-sized bins as shown in the figure. In each trial, the saccadic response was to one of two peripheral targets presented following fixation of a central target for 500-800 ms. This saccade direction was then discretized into the eight bins as described above. Each trial concluded 400-500 ms following the saccadic eye movement. The period between the start of a trial labeled Start and the Go cue is denoted as the Start-Go period, and the period between the Go cue and the end of the trial labeled End is denoted as the Go-End period.

#### 2.5.1. Data preprocessing

During task performance, extracellular potentials were acquired by a 32-electrode microdrive (Gray Matter Research) [47] and sampled at 30 kHz. Eye position was continuously tracked by an infrared eye-tracking system (ISCAN, USA) and sampled at 120 Hz. We extracted LFP power features after common average referencing from six frequency bands: delta–alpha (2-12 Hz), beta 1 (12-24 Hz), beta 2 (24-34 Hz), gamma 1 (34-55 Hz), gamma 2 (65-95 Hz), and gamma 3 (130-170 Hz) [6,8] with short-time Fourier transform estimates from 100 ms causal moving windows every 40 ms. We extracted spiking activity by band-pass filtering the raw neural signals between 300-6600 Hz and detecting the threshold crossings 3.5 standard deviations below the signal mean. We then downsampled the spiking activity to 250 Hz as there were rarely more than 1 spike per time bin. The resulting data thus had 4 ms Δ time-bins while the field features were observed at a slower 40 ms time-scale. We analyzed data from eight experimental sessions. These sessions contained a combined 3411 trials.

We analyzed each session individually with a fivefold cross-validation. For each training set and test set pair, we selected the top 40 LFP power features and the top 20 spiking channels based on saccade direction classification in the training set. Specifically, for each individual feature/channel and in the training set, we used linear discriminant analysis (LDA) with an inner four-fold cross-validation scheme to classify saccade direction and selected the features/channels with the highest average classification accuracy – note this selection was purely based on the training set. In the training set, the LDA classifier either used the average value of the LFP power feature or the average spiking rate over the Go-End regime in a one-versus-rest scheme, i.e. classify the correct direction among the 8 possible directions [48]. We use this same LDA classification to decode the brain states in the test set within cross-validation as detailed in Section 2.5.3.

#### 2.5.2. Model training

The behavioral task featured a switch between two regimes, which we term the Start-Go and the Go-End regimes, respectively. Thus, in the training set, we label these periods within each trial. We then assume we do not know the labels in the test set and when performing regime and brain state decoding. Having regime labels in the training set, we use an expectation-maximization (EM) learning algorithm for multiscale observations [14] to learn the dynamics equation parameters per regime and the observation equation parameters across regimes. We take the dimension of the brain state as *d* = 10 based on common values used in prior works [6, 8]. We then learn the regime state parameters using the regime labels concatenated over trials and maximumlikelihood techniques, and assume the initial regime state distribution is uniform for fairness.

#### 2.5.3. Switching multiscale filter (SMF) evaluation and performance metrics

We then apply SMF to the observations in the test sets to show that it can decode regimes and fuse information from multiscale observations. We quantify the regime state decoding using the accuracy metric in (48) to calculate how well the estimated regime states match the withheld regime labels. To assess the quality of brain state decoding, we see how well the decoded state averaged over the Go-End period could classify the direction of the saccadic eye movement response using LDA – note that the ground-truth brain state itself is latent and thus we quantify quality based on observed saccade directions. We do so in a one-versus-rest scheme [48] where for each possible direction we classify which of the saccades in the test set were in that direction to get an AUC metric, and then repeat this for each of the 8 directions and average the AUC scores.

## 3. Results

We show in both Monte Carlo simulations and in neural activity recorded from prefrontal cortex during behavior that the developed inference methods are capable of estimating both brain and regime states and fusing information from both Gaussian and point process observations. We use the Wilcoxon signed-rank test for all statistical comparisons and control for false discovery rate using the Benjamini-Hochberg Procedure [49]. Details on simulation settings for each section can be found in the appendix. Results figures with statistical tests performed contain p-values in their captions.

### 3.1. Switching inference improves estimation by tracking regime-dependent non-stationarity in simulations

We first find in simulations that the developed switching inference methods, SMF and SMS, significantly outperform their stationary counterparts across all swept simulation settings (Figure 4; *p* = 3.89 × 10^−8^). Using multiscale observations, the median reduction in error (NRMSE) when using the switching methods over stationary ones is 23.4% and could be as large as 34.2%. This reduction in error is due to the switching methods properly tracking regime-dependent non-stationarity as can be seen in Figure 4(a), where the median accuracy of regime state estimation exceeds 0.9 for the base simulation settings. We also find that when we also switch the eigenbasis as described in Methods, we get similar significant reductions in error of 25.0% and as large as 36.6% (*p* ≤ 3.89 × 10^−8^). Further, the benefit of switching methods increases as the number of regimes increase as seen in Figure 4(c), due to the stationary methods having to generalize for all regimes with only a single set of stationary parameters. Also, as seen in Figure 4(d), the performance of the switching methods improves as the dwell time increases as longer dwell times allow for longer periods of observations for regime identification. Finally, as seen in Figure 4(b), we find that the benefit of switching methods over stationary methods stays consistent even as the brain state dimension increases, which increases the difficulty of estimation since the same number of observation channels must encode increasing amounts of brain state information.

**Figure 4.**
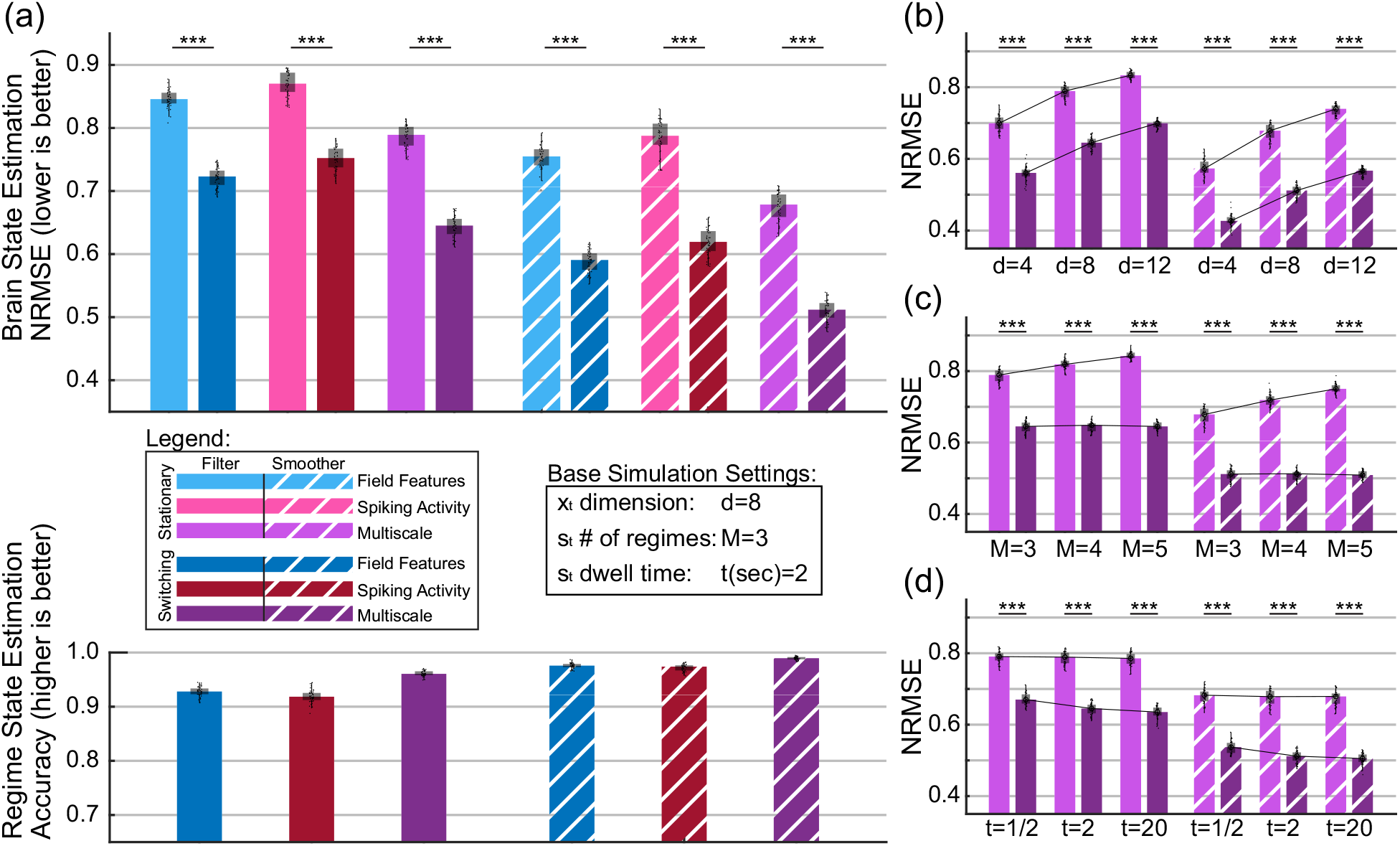
SMF and SMS improve estimation by tracking regime-dependent non-stationarity in simulations. (a) Brain state estimation comparison between stationary and switching inference methods across observation modalities: only field features, only spiking activity, and multiscale. Bars indicate the median NRMSE metric, box edges indicate the 25th and 75th percentiles, and whiskers indicate minimum and maximum values excluding outliers that exceed 1.5 times the interquartile distance. Base simulation settings are used in simulations throughout the figure unless otherwise overwritten by settings listed underneath bars. Asterisks indicate statistical significance with \*\*\**p* < 0.0005. (b) Comparison between multiscale stationary and multiscale switching inference NRMSE as brain state dimension sizes increase. Trend lines connect at the median. (c) Same as (b) but when the number of regimes *M* increases. (d) Same as (b) but when the dwell time *t* increases. For each statistical test, *n* = 40 and *p* = 3.89 × 10^−8^.

We then find that even with limited differences between regimes, SMF can still meaningfully detect the changes in regime. Specifically, to simulate limited changes, we use only spiking observations with 2 regimes and restrict what can switch to only the maximum firing rate as illustrated in Figure 5(a) such that the firing rate under the second regime is a fixed percentage smaller than that under the first regime, which we refer to as the percent difference between the regimes. This results in median errors in brain state estimation from a stationary filter that are within 0.03 NRMSE of those from a switching filter – thus this particular simulation setup generates only limited differences between regimes as intended. This simulation scenario was motivated by prior evidence [26] of similar switches in firing rates in spiking data recorded from the medial entorhinal cortex of mice in a random foraging task. Even with such limited differences between regimes, SMF could still achieve meaningful regime estimation accuracy. Indeed, when the percent difference in firing rate between regimes is greater than 40%, median accuracy values are greater than 0.8 as seen in Figure 5. Thus, SMF can yield useful estimates of the regime state even when differences between regimes are limited.

**Figure 5.**
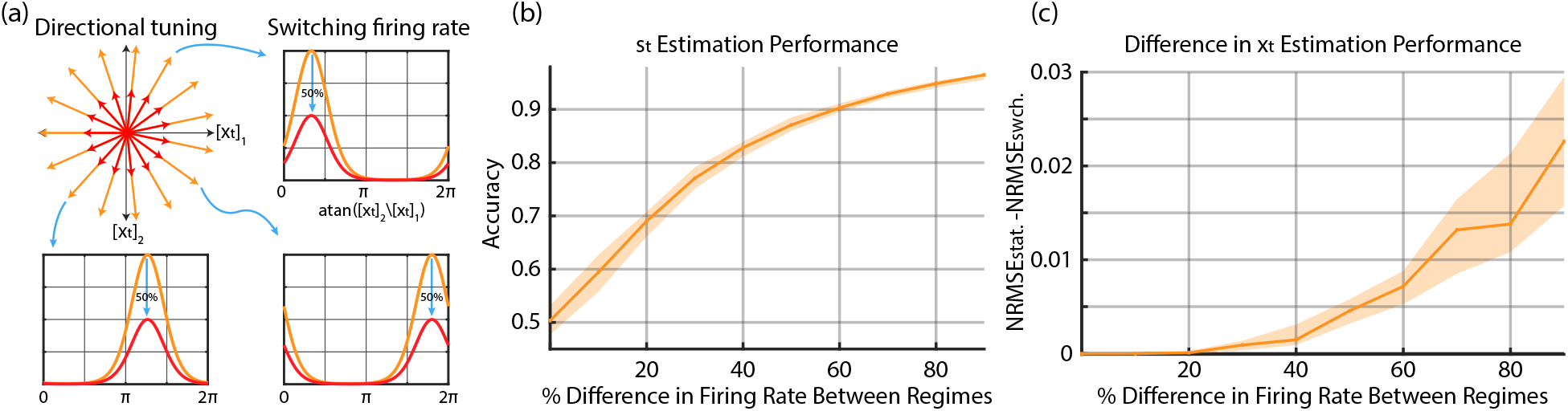
SMF can detect changes in regime corresponding to switches in even just the magnitude of spike firing rate in simulations. We set the dimension of the brain state **x**_*t*_ to 2, the number of regimes to 2, and the number of neurons to 15. (a) Illustration of how the magnitude of firing rates switch for the two regimes: orange and red. Top left shows the directional tuning of neurons, that is the **x**_*t*_ vector directions corresponding to maximum firing rate for all neurons, with each neuron associated with a different vector. The orange and red correspond to the two different regimes for each neuron, i.e., the two different maximum firing rates. Right and bottom show the firing rates for example individual neurons at the two regimes at a given **x**_*t*_ magnitude. Spike parameters are chosen for the first regime and then scaled down for the second such that the ratio of firing rates between the two is a specified percentage for all **x**_*t*_. We refer to this percentage as the percent difference between the regimes. (b) Regime state estimation as the percent difference in firing rate between regimes increases. Percent differences are chosen at 10 percent intervals between 0 and 90 with 40 systems simulated for each difference. Center line shows the median while shaded areas show 25th and 75th percentiles. (c) Difference in error (NRMSE) between stationary and switching methods as percent difference in firing rate increases. Positive values indicate better performance with SMF. We find significant differences between stationary and switching at percent differences ≥ 30.

### 3.2. Estimation performance improves by combining multiscale observations

We find that the new multiscale switching methods can robustly fuse information from Gaussian and point process observations and improve estimation performance. We sweep various settings such as maximum firing rate, field feature SNR, field feature sampling frequency, and regime state dwell time as seen in Figure 6. Across all settings, SMF and SMS get significantly better brain and regime state estimation when provided multiscale observations compared to that when provided only single-scale observations (*p* ≤ 3.12 × 10^−7^; Figure 6). Furthermore, we find that even if estimation performance with only field features is considerably worse than that with only spiking activity, or vice-versa, SMF and SMS can still yield improved performance with multiscale observations. This suggests that as long as there is some information in the worse performing single-scale modality, the multiscale switching methods can still fuse it with the better modality to achieve improved performance.

**Figure 6.**
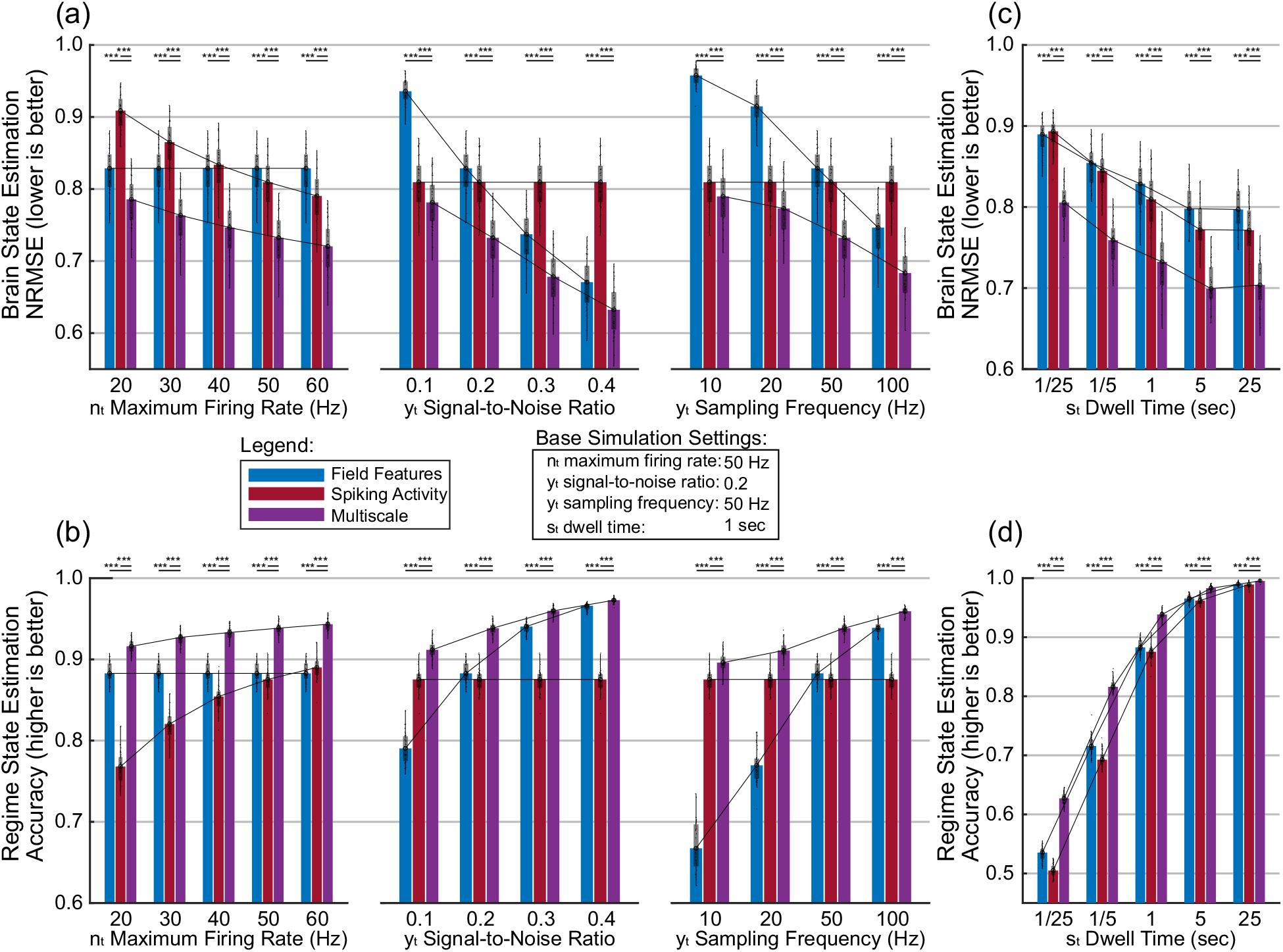
Switching methods improve estimation by combining multiscale observations in simulations. Bar, box, whisker, outlier, and asterisk conventions are the same as in Figure 4. Base simulation settings in box are used throughout the figure unless otherwise overwritten by individual settings listed underneath bars. (a) Brain state estimation performance across observation modalities under various observation parameter settings. Performance with multiscale observations is compared to that with only field features **y**_*t*_ and that with only spiking activity **n**_*t*_. For spike parameters, different maximum firing rates are simulated. For field feature parameters, different signal-to-noise ratios and different sampling frequency are separately simulated. (b) Regime state estimation performance across observation modalities under the same observation parameter settings used in (a). The number of regimes in these simulations is 3 and thus chance accuracy is 1/3. (c) Brain state estimation as regime state dwell time increases. (d) Regime state estimation as regime state dwell time increases similar to (c). For each statistical test, *n* = 40 and *p* ≤ 3.12 × 10^−7^.

### 3.3. Inference with switching multiscale smoother (SMS) is reliable across conditions

Our smoother is different from prior methods in two aspects. First, prior switching methods were for single-scale observations whereas our methods allow for multiscale observations. Second, even considering the smoothing inference step alone, we make a different approximation compared to prior methods – Kim’s method [31] and Expectation Correction (EC) [44] – as described in Equation (38). We make this new approximation to address the challenge of reliable performance across different simulation scenarios, which we refer to as generalizability. In particular, we find that, even with single-scale observations, while prior methods are accurate in some simulation scenarios, they can lead to unreliable smoothing under other simulation scenarios. Here, we find that our new switching smoother – whether multiscale or single-scale – can address this challenge of reliable performance across diverse settings and thus generalize well.

Under a setting tested from Section 3.2 with additionally switching eigenbases, we find that prior smoothing methods – Kim’s method [31] and EC [44] – yield significantly worse error compared to SMS across all observation types whether single-scale or multiscale (*p* ≤ 9.83 × 10^−7^) with settings and plots in Figure 7(e). We note that this is the case even though the settings are not extreme in value, with all systems being numerically bounded. Furthermore, we see that prior smoothing methods can yield worse error than even their filtering counterparts. For example, for the spiking observation modality, this occurs for more than 50% of the systems tested (25/40 systems for Kim’s method, 29/40 for EC) as seen in Figure 7(d). In comparison, SMS reduces error and increases accuracy compared to its filtering counterpart (SMF) across all observation types, whether single-scale or multiscale, and for every system tested not only under these settings but also for all settings in Section 3.2. Also importantly, this improvement in generalizability across diverse settings comes at only a slight price in performance in some specific settings. In particular, for settings from Section 3.2 which have stationary eigenbases and thus yield systems that are easier to track for prior methods, SMS only has a 4.0 × 10^−4^ and 3.16 × 10^−4^ median increase in NRMSE compared to Kim’s method and EC respectively. Further, SMS’s accuracy in regime estimation in these settings is comparable to Kim’s methods and only 4.1 × 10^−4^ lower compared with EC. A subset of these comparisons shown on the right side of Figure 7. These results show that SMS smoother reliably improves estimation compared to the SMF filter and can robustly generalize across diverse conditions unlike prior methods.

**Figure 7.**
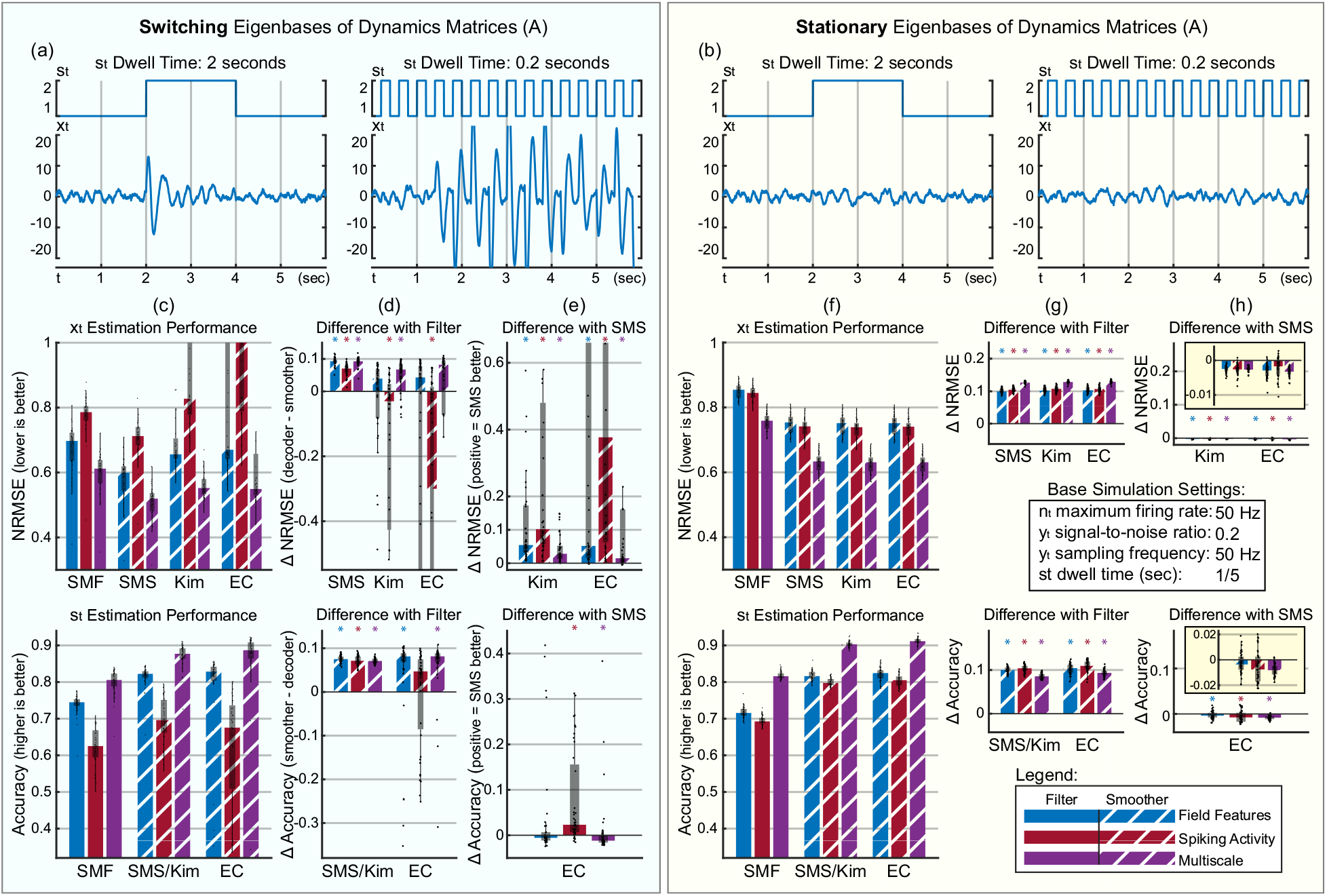
SMS smoothing reliably improves estimation compared to filtering across diverse settings, thus enabling generalizability unlike prior smoothing methods. Bar, box, whisker, and outlier conventions are the same as in Figure 4. (a) Example traces when the eigenbases, comprised of the eigenvectors of the dynamics matrix **A**, can switch across regimes. In this setting, differences in eigenbases across regimes can lead to transient responses on regime switch that can compound for shorter dwell times. SMS markedly outperforms prior smoothing methods when eigenbases are switching as seen in (c–e). (b) Example regime and brain state traces when the eigenbases are stationary across regimes. Note that in this setting in contrast to the setting in (a), even with changes in dwell time, the overall magnitude of the brain state signal remains relatively the same. (c) Brain (*top*) and regime (*bottom*) state estimation using the filter SMF and the three smoothing methods, SMS, Kim’s method, and EC, in the case with switching eigenbases and a dwell time of 0.2 seconds. SMS outperforms Kim’s method and EC in smoothing. Note that in the bottom panel, SMS and Kim’s method use the same regime state smoothing equations and thus yield the same accuracy (this is not the case for the brain state in the top panel though). (d) Pair-wise differences in brain and regime state estimation between the filter SMF and the three smoothing methods. Positive values indicate better performance using a smoother. (e) Pair-wise differences in estimation between SMS and other smoothing methods. Positive values indicate better performance using SMS. SMS outperforms prior smoothing methods. (f–h) Equivalent plots and conventions to (c–e) applied to systems with stationary eigenbases. Insets in (h) show zoomed in versions of their respective underlying plots as SMS performs very closely to prior smoothing methods even in this case while allowing for generalizability to other settings as seen in panels (c–e). Simulation settings used through this figure listed in Base Simulation Settings box. Asterisks show whether significant or not (*n*=40, \**p* ≤ 0.05).

### 3.4. Switching multiscale filter (SMF) detects task-related regimes in spike-field NHP data

We then find that using the developed SMF method with real spike-field neural observations, we can causally and accurately detect regimes related to the saccade task performed by a monkey (Figure 8). We train and test the switching multiscale dynamical system (SMDS) models in cross-validation (see Section 2.5.2). We find that the switching multiscale filter (SMF) can successfully decode whether multiscale spike-LFP activity belong to the Start-Go or the Go-End regimes of the task with a median accuracy of 0.808 as seen in Figure 8(d). This estimation is done based on SMF computing the conditional probabilities of the two task-related regimes, and then picking the one with the higher conditional probability (i.e. Start-Go regime and Go-End regime, see Section 2.5.2).

**Figure 8.**
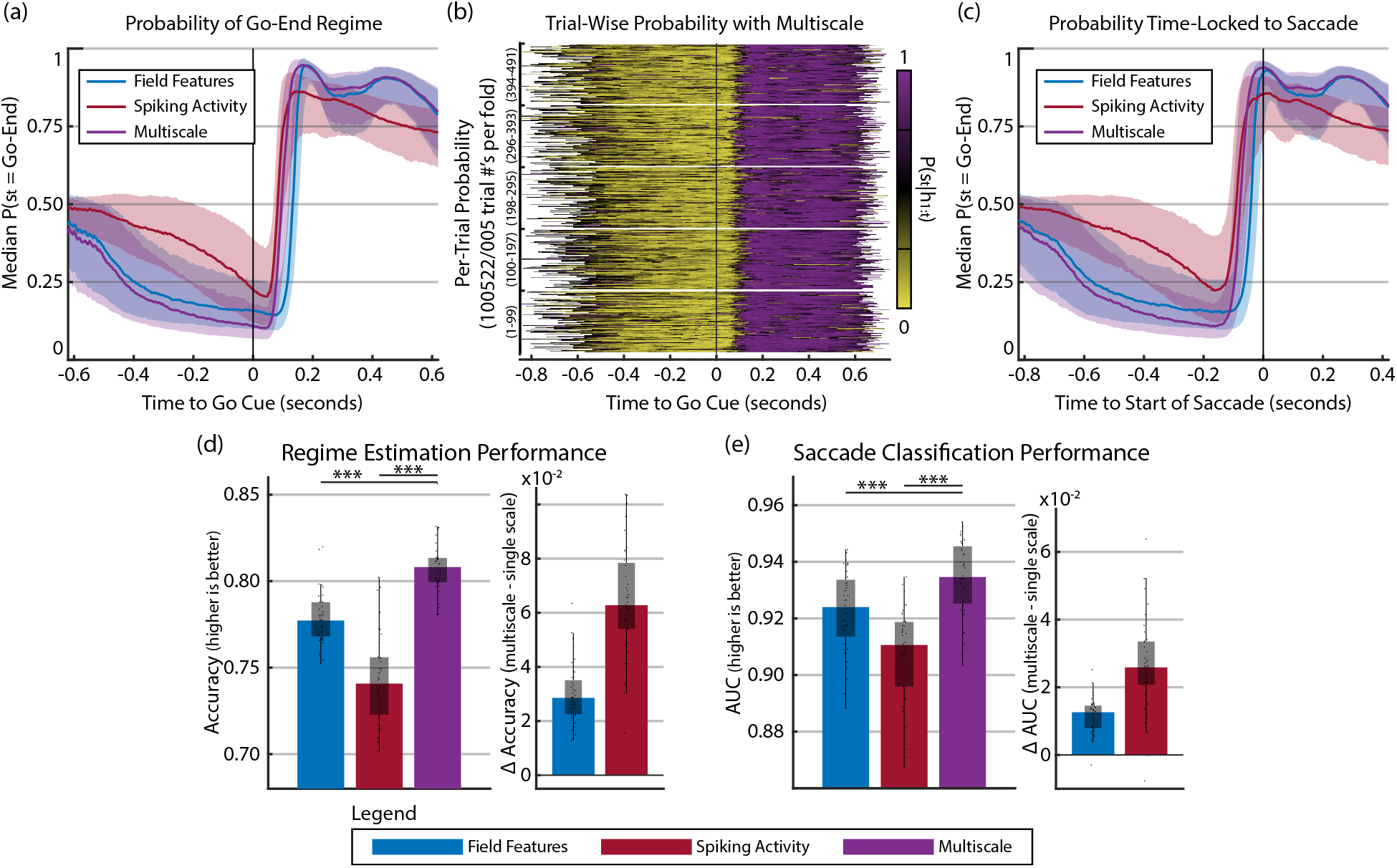
The switching multiscale filter (SMF) method can perform real-time multiscale decoding of task-related brain and regime states in data recordings from a non-human primate and combine information from spike and LFP scales. (a) Resulting conditional probability of the Go-End regime from SMF time-locked to the Go cue. Center lines represent the median across trials from all sessions while shaded areas represent 25th and 75th percentiles. Perfect decoding would be a step signal 0 valued prior to Go and 1 valued after Go. (b) Individual trials from the five test folds in one example session with colors indicating the conditional probability of the Go-End regime according to the color bar. Trials are time-locked to the Go cue and from SMF being applied to multiscale spike-LFP observations. (c) The same probability traces as in (a) but time-locked to detected saccade onset at time 0. (d) Performance in decoding Start-Go and Go-End regimes across observation modalities. Bar, box, whisker, outlier, and asterisk conventions are the same as in Figure 4. Right side shows pair-wise differences between performance with multiscale to that with LFP (blue) and spiking (red) with positive values indicating better multiscale performance. (e) Performance in classifying saccade direction using the mean of decoded brain state signal from Go-End regime (see Methods) across observation modalities with pairwise differences on right as in (d). *n* = 40 for all statistical tests.

Furthermore, the computed conditional probability of the Go-End regime for even individual trials clearly and rapidly increase once the saccade targets appear at the Go cue as seen in Figure 8(b) and Figure A1. The median time-locked trace of these probabilities are shown in Figure 8(a). We note that this increase in probability occurs prior to saccade initiation as seen in Figure 8(c), suggesting that SMF can detect changes in multiscale neural dynamics prior to any detectable movement.

We also find that not only could we classify regimes related to the behavioral task, but also we could use the decoded brain state from SMF to accurately classify the direction of the saccade with a median AUC of 0.935 as seen in Figure 8(e). This AUC is significantly better than the AUC from stationary models (*n* = 40, *p* = 0.00838). This result suggests that SMF can accurately find task-relevant brain and regime states from multiscale observations.

Finally, consistent with our results in Section 3.2, we find that regime and saccade direction classification performance is significantly better using multiscale spike-LFP observations compared to using single-scale observations (whether spike or LFP) as seen throughout Figure 8. Overall, we find that the performance of SMF on spike-field activity in prefrontal cortex yields results that are consistent with results based on simulated data. The results show the potential of the developed switching multiscale inference methods for investigations of neural activity with switching dynamics and/or encoding and for brain-machine interface applications.

## 4. Discussion

We developed the switching multiscale dynamical system (SMDS) model that generalizes the common linear dynamical system (LDS) and developed companion filtering/decoding and smoothing algorithms. Our model incorporates two advances: it augments the LDS by modeling an additional binary-valued point process observation modality; it further relaxes the stationarity assumption of the resulting multiscale dynamical system model with the inclusion of an additional regime state that governs the model parameters. We also designed a real-time decoding algorithm for the SMDS model termed the switching multiscale filter (SMF), which simultaneously estimates regime and brain states from spike-field activity while maintaining a tractable computational cost. We then developed a fixed-interval smoothing algorithm termed the switching multiscale smoother (SMS), which can further increase estimation accuracy compared to SMF by utilizing the entire length of data across a broad range of model settings when causal estimation is not needed.

To validate the methods, first, we conducted extensive numerical simulations and compared with stationary multiscale methods or switching single-scale models (e.g. switching LDS) that use only one observation modality. Doing so, we showed that our framework improves estimation for spike-field activity with regime-dependent dynamics by successfully leveraging these different observation modalities simultaneously to both track regime-dependent non-stationarity and decode brain states. Second, we tested our methods under extensive simulation settings to show their robustness and characterize how various settings can impact estimation performance. Finally, we used experimental recordings of neural activity in the prefrontal cortex of one non-human primate and validated that our methods can perform real-time decoding and simultaneously estimate task-related brain and regime states from multiscale spike-LFP activity.

The presented modeling and inference framework has the potential to improve brain-machine interfaces (BMIs) for decoding and modulation of brain states, for example for the purpose of restoring lost motor function in disabled patients [1, 2, 5]. As neurotechnologies are developed for long-term use, inference algorithms must develop in parallel to become more appropriate for naturalistic multi-regime scenarios where there could exist multiple tasks with different strategies and/or complex tasks with multiple discrete stages/regimes. In addition, higher level cognitive states such as a subject’s level of stress, engagement, or attention can induce multiple regimes in the task by impacting the strategy and thus the dynamics of behavior and neural activity [23–25]. Indeed, not only can the temporal dynamics of brain states change, how they are encoded in neural observations can also switch due to external or internal cues [26]. Regime-detection methods are an answer to solving these challenges. Further, as technology advances, BMIs will have access to simultaneous spike-field recordings and thus new methods are needed to optimally combine these modalities for improved performance and robustness. We thus provide a novel regime-detection method that can incorporate multiscale spike-field measurements using combined Gaussian and point process observations.

Our work extends traditional regime-detection algorithms for BMIs. Prior work has used hidden-Markov models (HMM’s) for task-related regime detection in parallel with a continuous state inference algorithm [50]. But in this case, the HMM does not influence the model used by the inference algorithm and so the inference algorithm does not directly consider the switching. Other prior work has addressed the case where a single but unknown regime is in effect over the entire course of a trial [51]. This is done by maintaining a finite number of separate filters each corresponding to one possible regime and, on a trial-by-trial basis, updating the likelihood of which regime is in effect for that entire trial [51]. However, this approach is designed for situations where only one regime is in effect over the entirety of a trial and not for situations where the regime can actually switch over the course of a trial or during longer naturalistic recordings that lack a controlled trial-based structure. Applying the approach of this prior work to the latter situations can lead to error in the filters accumulating since the separate filters do not interact with each other; this error accumulation has been shown in the classical switching LDS systems [30]. Finally, other prior work has considered situations where the regimes can switch, but the algorithm only takes single-scale Gaussian observations and is not applicable to the multiscale case as it does not perform inference on simultaneous Gaussian and point process observations [27]. In summary, the modeling and inference framework developed here has the potential to improve BMIs in more naturalistic scenarios and to further take advantage of multiscale measurements to enhance performance and robustness.

The presented framework could also become a valuable tool in estimating mental states such as mood that are represented across distributed brain regions, such that they can be used as feedback to personalize therapies for mental disorders such as deep brain stimulation [5, 32, 33, 36]. Recent work has shown successful decoding of mood states in individuals from multiple cortico-limbic regions [5, 32]. These regions are often involved in various functions and internal states such as attention, salience, and cognitive control [5], which can affect the dynamics and/or encoding of a given mental state by imposing various regimes. By tracking these regime-dependencies, our framework could thus improve the decoding of mental states.

In this paper, we focused on developing the SMDS model and companion inference methods and performed validation with extensive simulations as well as NHP data. The extensive simulations allowed us to test the generality of our methods across a wide range of conditions such as regime-switching frequency and model parameters. They also provided the opportunity to elucidate how various conditions can impact estimation performance and in what situations our methods yield greater advantages over existing methods by comparing to known groundtruths. Prior works have similarly shown that simulations are a valuable tool for developing new algorithms for switching regime systems [44, 52, 53] and neurotechnologies in general [6, 7, 14, 19, 20, 35, 41, 54–60].

Motivated by prior work that show spiking activity and field features could be approximately conditionally independent conditioned on the unobserved brain states [7, 8], we made a similar assumption in our inference algorithms. Consistently, we showed in the experimental data from prefrontal cortex that our models and inference methods were successful in estimating both task-related regimes as well as saccade direction from spike-field activity. To develop neurotechnologies, this assumption also yields computational advantages in terms of limiting the number of parameters to learn and thus reducing the amount of training data and improving model accuracy and generalizability. Nevertheless, future work that further models any potential conditional dependencies could provide a scientific tool to explore the functional connectivity between spiking activity and field features [59,61–63]. However, any such future work will have to address the issue of having substantially larger number of parameters that need to be fitted with limited training data, which would make the models prone to overfitting. Thus, new learning methods may be required to address this issue.

In the experimental data from prefrontal cortex, we could label the regimes based on external task cues in the training set, and thus learn our models parameters supervised with respect to the regime state. However, in other experimental paradigms, task or neural regimes may be due to internal rather than external cues. Thus, future work can develop unsupervised learning algorithms for regime-dependent models in which regime labels may not be available in the training set. Recent work has developed such unsupervised methods for single-scale observations with Bayesian and variational approximation methods [52, 53]. Future work should develop such methods for multiscale observations and explore the new unique challenges that arise from modeling these multiple scales simultaneously. Our multiscale inference methods can provide a key computational element for developing such unsupervised multiscale learning algorithms.

## 5. Acknowledgments

This work was supported in part by Army Research Office (ARO) under contract W911NF-16-1-0368 as part of the collaboration between US DOD, UK MOD and UK Engineering and Physical Research Council (EPSRC) under the Multidisciplinary University Research Initiative (MURI). The authors thank David Markowitz who designed the non-human primate experiments and collected the non-human primate data in the Pesaran Lab.

## Appendix A. Details on simulation settings

For the main set of comparisons in Section 3.1, we sweep all combinations of **x**_*t*_ dimension *d* = [4,8,12], number of regimes *M* = [3,4,5], and regime dwell times *t_dwell_* = [0.2, 2, 20] seconds. For observation equation parameters, we set **y**_t_ SNR randomly to between [0.2,0.24], **y**_*t*_ sampling frequency to 50Hz, and the maximum firing rate for **n**_*t*_ to 40Hz. For comparisons where we allow for switching eigenbases, we set observation equation parameters to be stationary and set *d* = 8, *M* = 3, the transient control term *σ* = 3, and swept *t_dwell_* = [0.2,2,20] seconds. For comparisons where only the magnitude of spiking activity parameters switch, we set *d* = 2, *M* = 2, *t_dwell_* = 2, and the number of neurons to 15.

For the main set of comparisons in Section 3.2, as base settings, we set *d* = 8, *M* = 3, *t_dwell_* = 1 second, maximum firing rate of **n**_*t*_ to 50Hz, **y**_*t*_ SNR to 0.2, and **y***t* sampling frequency to 50Hz. We then individually sweep values for the following settings while holding the remaining at the base settings: maximum firing rate of **n**_*t*_, **y**_*t*_ SNR, **y**_*t*_ sampling frequency, and *t_dwell_*. For maximum firing rate of **n**_*t*_, we swept frequencies 20 – 60Hz in 10Hz increments. For **y**_*t*_ SNR, we swept values 0.1 – 0.4 in 0.1 increments. For **y***t* sampling frequency, we swept frequencies [10, 20, 50,100]Hz. For *t_dwell_*, we swept times [1/25,1/5,1, 5, 25] seconds.

For Section 3.3 where we allow eigenbases to switch, we set *d* = 8, *σ* = 3, *M* = 3, *t_dwell_* = 0.2 seconds, maximum firing rate of **n**_*t*_ to 50Hz, **y**_*t*_ SNR to 0.2, and **y**_*t*_ sampling frequency to 50Hz.

**Figure A1.**
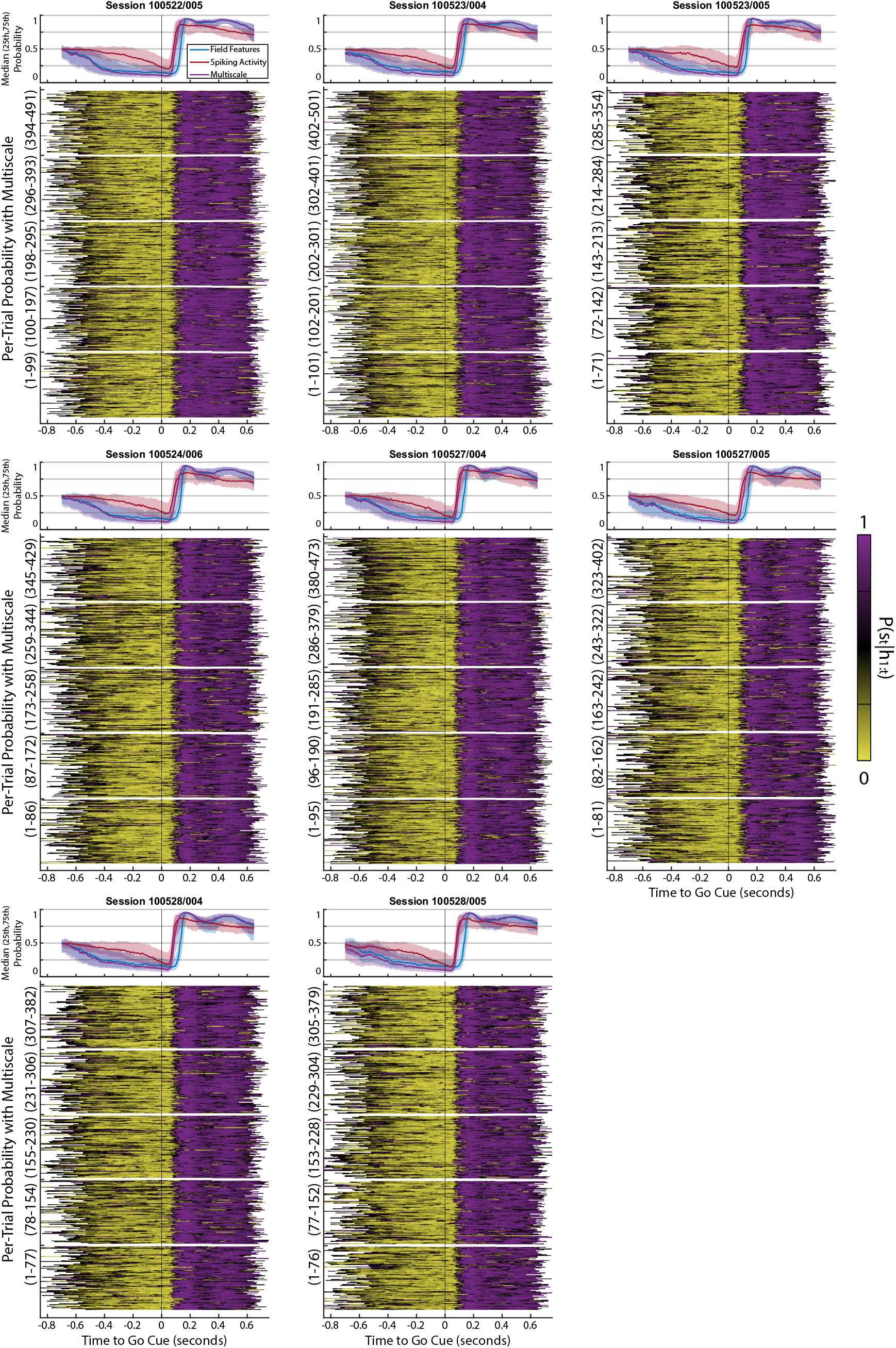
Conditional probabilities of the Go-End regime from each individual recording session. *Top panel per session:* Conditional probability that a time point is from the ‘Go-End’ regime, time-locked to the Go cue across observation modalities. Center lines represent the median across trials from that session while shaded areas represent the 25th and 75th percentiles. *Bottom panel per session:* Individual trials from the five test folds from that session with colors indicating the conditional probability of the Go-End regime according to the color bar. Trials are time-locked to the Go cue and from SMF being applied to multiscale spike-LFP activity.

## Notes

### Competing Interest Statement

The authors have declared no competing interest.

